# Attenuated replication and damaging effects of SARS-CoV-2 Omicron variants in an intestinal epithelial barrier model

**DOI:** 10.1101/2024.02.28.582510

**Authors:** Meta Volcic, Rayhane Nchioua, Chiara Pastorio, Fabian Zech, Clarissa Read, Paul Walther, Frank Kirchhoff

## Abstract

Many COVID-19 patients suffer from gastrointestinal symptoms and impaired intestinal barrier function may play a key role in Long COVID. Despite its importance, the impact of SARS-CoV-2 on intestinal epithelia is poorly understood. To address this, we established an intestinal barrier model integrating epithelial Caco-2 cells, mucus-secreting HT29 cells and human Raji cells. This gut epithelial model allows efficient differentiation of Caco-2 cells into microfold-like cells, faithfully mimics intestinal barrier function, and is highly permissive to SARS-CoV-2 infection. Early strains of SARS-CoV-2 and the Delta variant replicated with high efficiency, severely disrupted barrier function, and depleted tight junction proteins, such as claudin-1, occludin and ZO-1. In comparison, Omicron subvariants also depleted ZO-1 from tight junctions but had fewer damaging effects on mucosal integrity and barrier function. Remdesivir and the TMPRSS2 inhibitor Camostat prevented SARS-CoV-2 replication and thus epithelial barrier damage, while the Cathepsin inhibitor E64d was ineffective. Our results support that SARS-CoV-2 disrupts intestinal barrier function but further suggest that circulating Omicron variants are less damaging than earlier viral strains.

## INTRODUCTION

SARS-CoV-2 is primarily transmitted through respiratory droplets and the primary targets of viral infection are cells of the sino-nasal airway epithelium (Ahn et al., 2021). Frequently, SARS-CoV-2 replication is limited to the upper respiratory tract and most infections remain asymptomatic or mild (Lamers and Haagmans, 2022; V’kovski et al., 2021). In some cases, however, the virus may spread more systemically to the lower respiratory tract as well as to different organs causing a broad range of severe and sometimes life-threatening symptoms (Gavriatopoulou et al., 2020; Song et al., 2021). In addition to respiratory complications, up to 30% of COVID-19 patients experience gastrointestinal symptoms, including diarrhea, abdominal discomfort, loss of appetite and vomiting (Al-Momani et al., n.d.; Hayashi et al., 2021; Zhong et al., 2020). Some COVID-19 patients even develop severe duodenitis associated with gastrointestinal bleeding requiring red blood cell transfusion (Cappell and Friedel, 2023; Eleftheriotis et al., 2023). Impaired intestinal barrier function in SARS-CoV-2 infection allows microbial and endotoxin translocation, which triggers inflammation and may lead to sepsis or contribute to chronic inflammation (Assimakopoulos et al., 2022; Eleftheriotis et al., 2023; Yamada et al., 2022). Altogether, accumulating evidence suggests that SARS-CoV-2 infection of the gastrointestinal tract plays a relevant role in COVID-19 (Jin et al., 2021; Scaldaferri et al., 2020; Zhang et al., 2021). Notably, gastrointestinal disorders may persist after the acute phase of COVID-19 and contribute to post-acute sequelae of SARS-CoV-2 infection known as Long-COVID (Xu et al., 2023).

It has been reported that angiotensin-converting enzyme 2 (ACE2), the primary receptor of SARS-CoV-2 (Hoffmann et al., 2020), is expressed at higher levels by intestinal cells compared to lung cells (Guimarães Sousa et al., 2022; Rahban et al., 2021). Thus, both, direct SARS-CoV-2 infection and inflammatory cytokines, may compromise gut barrier integrity in COVID-19 patients (Jiao et al., 2021; Kariyawasam et al., 2021). Immunohistochemical analysis of gut biopsies from COVID-19 patients stained positive for the SARS-CoV-2 Spike protein and *in situ* hybridization analysis further support active viral replication (Neuberger et al., 2022). Previous studies suggest that SARS-CoV-2 affects various aspects of gut immune and barrier function and show that the integrity of the intestinal barrier is significantly impaired in a notable portion of COVID-19 patients (Eleftheriotis et al., 2023; Farsi et al., 2022; Tsounis et al., 2023; Yamada et al., 2022; Zuo et al., 2020). Despite its importance in the pathogenesis of COVID-19, the impact of SARS-CoV-2 infection on intestinal barrier function is poorly explored. Here, we examined the replication potential of various SARS-CoV-2 strains, *i.e.* French, Netherland, Delta and Omicron BA.1, BA.2, BA.5 and XBB1.5, in intestinal epithelia and their effect on barrier permeability, cell integrity, and tight junction proteins. To achieve this, we optimized a Caco-2 and HT29-MTX co-culture *in vitro* cell model for the permeability and function of the intestinal epithelium (Béduneau et al., 2014; Mahler et al., 2009; Pan et al., 2015) to study SARS-CoV-2 infection. Co-culture of intestinal Caco-2 enterocytes and HT29-MTX goblet cells resulted in an epithelial layer with strong cellular polarity, tight junctions and the presence of a thick mucus layer. The co-cultures expressed high levels of ACE2, as well as the transmembrane serine protease 2 (TMPRSS2) that activates the viral Spike protein, and were highly permissive to SARS-CoV-2 replication after apical viral exposure. Notably, differentiation of Caco-2 cells to M (microfold)-like cells by co-culture with Raji cells (Masuda et al., 2011) increased susceptibility for productive viral infection from the basolateral side. Early SARS-CoV-2 strains and the Delta VOC depleted tight junction proteins and impaired barrier function more severely than Omicron subvariants.

## RESULTS

### A triple co-culture model for intestinal SARS-CoV-2 infection

To investigate the ability of SARS-CoV-2 to infect intestinal epithelial cells and to affect gut permeability, we established a triple co-culture model comprising enterocytes, goblet cells and M cells similar to those used to study the transport of bacteria, proteins or drugs across mucosal barriers (Béduneau et al., 2014; Mahler et al., 2009; Pan et al., 2015). We used a trans-well system where human colon carcinoma Caco-2 cells, presenting enterocytes, and HT29-MTX cells, presenting mucus-secreting goblet cells, were seeded onto the apical side of the membrane (Figure 1A). In the initial experiments, we seeded either Caco-2 or HT29 cells alone, or two different mixtures of both cell types to achieve appropriate trans-epithelial electrical resistance (TEER) (between 300-450 Ω for 7:3 Caco-2 and HT29 mixed monolayer), an optimal ratio between enterocytes and goblet cells, and a proper mucus distribution (Figure 1B). During the first 14 days of culture confluent monolayers with cells expressing apicobasal polarity are formed. As tight junctions developed, TEER values increased over time, reaching the highest values (up to 800 Ω) in the Caco-2 single-cell monolayer, followed by the 9:1 and 7:3 mixtures of double-cell type monolayers (Figure 1B). The HT29 single-cell monolayer showed the lowest TEER values (<100 Ω) since goblet cells alone form less tight junctions. At day 14, human B lymphoma Raji cells were added to the basolateral compartment for an additional 5 days (Figure 1A). Comparison of co-cultures otherwise kept under identical conditions showed that Raji B cells triggered differentiation of Caco-2 cells into M-like cells, resulting in a reduction in TEER (Figure 1C) and alkaline phosphatase activity (Figure 1D). The presence of M cells was further confirmed by staining actin filaments with Phalloidine, revealing a thinning of the apical layer due to the loss of microvilli in M-like cells (Figure 1E). Additionally, scanning electron microscopy (SEM) confirmed the presence of M-like cells, where microvilli were sparser and less dense than in Caco-2 cells (Figure 1F). One characteristic of the intestine is the presence of mucus produced by goblet cells (Kim and Ho, 2010). In this model, HT29-MTX mucus-secreting differentiated goblet cells were used (Gagnon et al., 2013). Alcian Blue staining was performed to evaluate the optimal ratio between enterocytes and goblet cells for producing a homogeneously distributed mucus layer. The HT29 single-cell monolayer demonstrated blue staining across the entire surface, while this was lacking in the Caco-2 single-cell monolayer (Figure 1G). The 9:1 Caco2-HT29 mixture resulted in sparse blue staining, whereas a more even distribution was observed for the 7:3 mixture. Thus, we used the 7:3 mixture of Caco-2 and HT29 cells in subsequent experiments.

**Figure 1:**
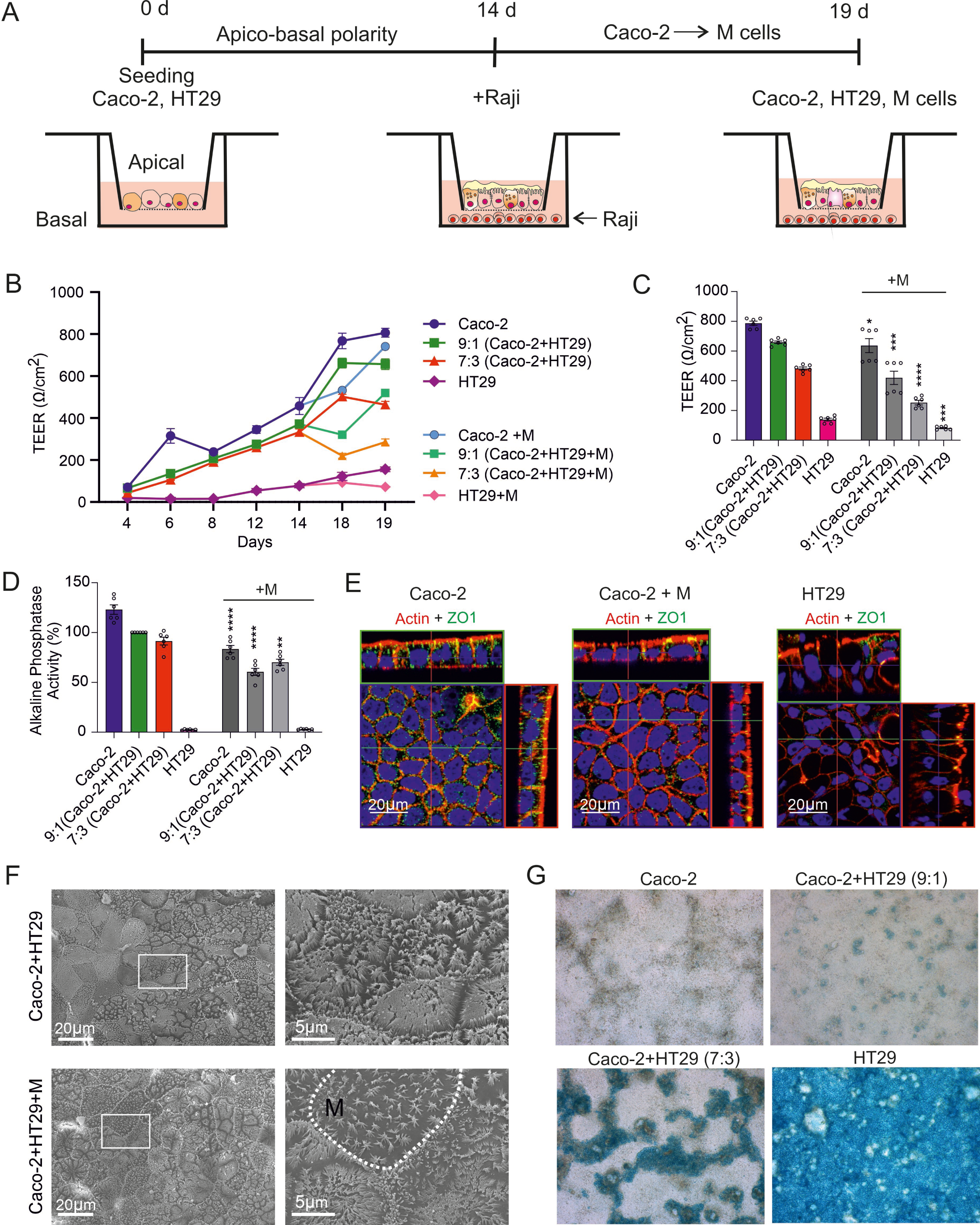
*In vitro* intestinal epithelial model. (**A**) Intestinal epithelial model. (**B**) Measurement of TEER of Caco-2, HT29 monolayer or combination of both cell types in different ration (9:1 and 7:3) in 19d time. (**C**) TEER values and (**D**) alkaline phosphatase assay on a day 19 of Caco-2, HT29 or mixture of both cell types in a ratio 9:1 or 7:3 with or without addition of Raji cells (+M). Caco-2+HT29 9:1 set to 100%, n=6, the bars represent the mean ± s.e.m.; \**p* = 0.0117, \*\**p* = 0.0016 and \*\*\**p* = 0.0004, **** *p*< 0.0001; unpaired *t*-test. (**E**) Immunofluorescence staining of beta-actin (red) and ZO-1 (green) Caco-2, Caco-2 + M and HT29 monolayer. (**F**) Scanning electron microscopy images of Caco-2 and Caco-2+M cellular monolayer confirming the presence of M cells with scarce microvilli formation. (**G**) Light microscopy images of Alcian blue mucus staining showing that HT29-MTX cells are producing mucus in single cell culture as well as in co-culture with Caco-2 cells. Mucus was not detected in Caco-2 single cell culture.

### SARS-CoV-2 replicates efficiently in the triple-cell intestinal model

To determine whether the intestinal epithelial model allows productive SARS-CoV-2 infection, the intestinal monolayer was exposed to the SARS-CoV-2 FR strain isolated in France in 2020 at a multiplicity of infection (MOI) of 0.1 for 6 hours (Figure 2A). Subsequently, the cells were washed to remove unbound virus, fresh medium was added, and cells were cultivated for another one to three days, followed by analyses for virus production and cell monolayer permeability. Increasingly high levels of SARS-CoV-2 RNA were detected in the supernatants of the epithelial cell cultures obtained at 1, 2 and 3 days post-infection (Figure 2B). Treatment with the RNA polymerase inhibitor Remdesivir reduced the levels of viral RNA to the detection limit. Western blot analyses confirmed efficient expression of the SARS-CoV-2 nucleocapsid (N) protein in the absence but not in the presence of Remdesivir (Figure 2C). SARS-CoV-2 FR infection disturbed the integrity and greatly increased the permeability of the cell monolayer (Figure 2D). In agreement with the high susceptibility of the intestinal model to SARS-CoV-2 replication, differentiated Caco-2/HT29 co-cultures (Intestinal Model) expressed high levels of ACE2 as well as TMPRSS2 (Figure 2E) (Hoffmann et al., 2020). Notably, ACE2 expression levels in differentiated Caco-2/HT29 co-cultures were about 80- and 8-fold higher compared to those detected in individual Caco-2 and HT29 cell cultures, respectively (Figure 2E). In addition, expression of TMPRSS2 was also increased in differentiated Caco-2/HT29 cultures (Figure 2E). It has been reported that SARS-CoV-2 infection down-regulates ACE2 in the respiratory epithelium (Perrotta et al., 2020). In agreement with this, we found that upon SARS-CoV-2 infection ACE2 expression in the intestinal monolayer model was reduced by ∼3-fold (Figure 2F). Next, we performed scanning electron microscopy (SEM) to investigate the impact of SARS CoV-2 on morphology of cellular micro villi. In uninfected samples we observed dense and healthy-looking microvilli, whereas in FR or Delta infected samples micro villi were sparser (Figures 2G, S1). Moreover, SEM revealed high numbers of SARS-CoV-2 particles attached to or released from the cellular villi (Figures 2G, S1), which agrees with previous data showing that ACE2 is expressed by intestinal villi (Lee et al., 2020). Altogether, the results demonstrated that differentiated Caco-2/HT29 co-cultures are highly susceptible to SARS-CoV-2 replication and that they are a useful model to examine viral effects on gastrointestinal barrier function.

**Figure 2.**
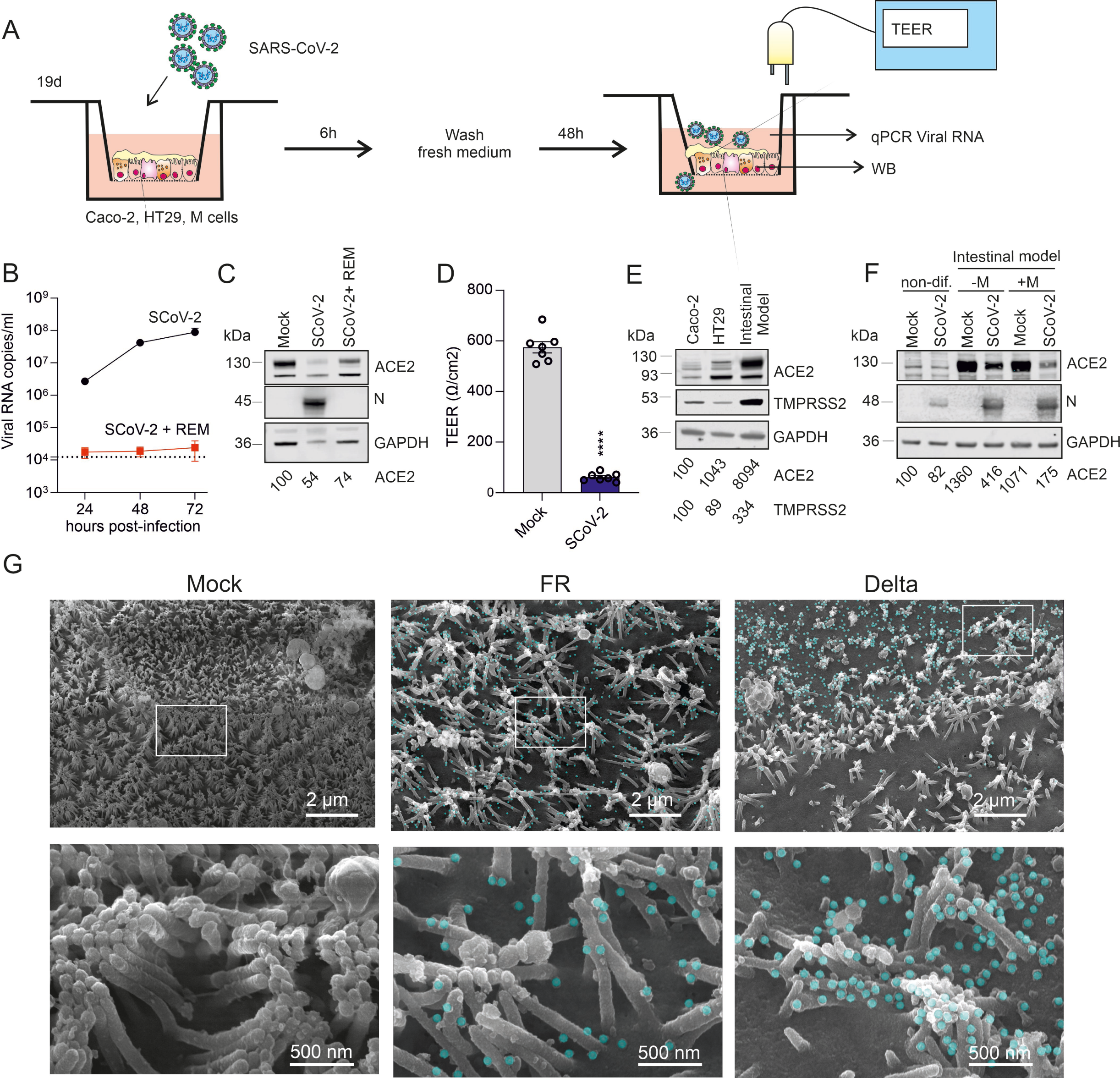
SARS CoV-2 infection of intestinal epithelial model. (**A**) Experimental set-up. On a day 19, Raji cells were removed, fresh medium containing SARS CoV-2 virus at MOI 0.1 was added into the apical compartment. Cells were incubated for 6h, when virus was washed away. After 48h cultivation, TEER was measured, supernatants and cells were taken for qPCR or WB analysis. (**B)** Viral RNA copies/ml 24h, 48h and 72h poi in supernatant from Intestinal epithelial model which was pretreated or not with Remdesivir and infected with FR SARS CoV-2. (**C**) WB presenting ACE2 and N expression in intestinal epithelial model which was mock infected, infected or pretreated with Remdesivir and infected with FR SARS CoV-2. Cells were collected 72h poi. (**D**) TEER measurement of intestinal epithelial model which was mock infected or infected with FR SARS CoV-2 strain. n=7, the bars represent the mean ± s.e.m.; **** *p*< 0.0001; unpaired *t*-test. (**E)** Westrn blot presenting expression of ACE2 and TMPRESS2 in Caco-2, HT29 non-differentiated cells and in intestinal model (differentiated cells). (**F**) Expression of ACE2 and N protein in non- and differentiated Caco-2 or Caco-2+M cells infected with FR SARS CoV-2 strain. (**G**) SEM images of intestinal model infected with FR, Delta SARS CoV-2 or mock treated. Green colored are viral particles detected by the use of program 3dmod with size criteria 70-100nm.

### M cells support basolateral SARS-CoV-2 infection of the intestinal epithelium

After establishment of the model, we examined the impact of M cells on the susceptibility of the intestinal epithelium to SARS-CoV-2 infection and on barrier function. M cells endocytose macromolecules, particles and microorganisms from the intestinal lumen and exocytose them to their basolateral membranes where T and B lymphocytes are present (Casteleyn et al., 2013). Thus, they play key roles in mucosal immunity and transcytosis. Differentiation of Caco-2 cells to M-like cells can be induced by co-culture with Raji cells and is relevant for nanoparticles uptake in related intestinal models (Cabellos et al., 2017). The gut has been suggested as an alternative entry site of SARS-CoV-2 and may be infected both directly or by systemic spread of the virus from the lung and other organs (Clerbaux et al., 2022; Guo et al., 2021). Thus, we challenged the gut epithelium model with the virus from both the top and the bottom. Infection from the apical side was associated with effective replication of the FR and Delta strains in both the presence and absence of M cells (Figure 3A). On average, the Omicron BA.1 variant produced about 4 orders of magnitude lower levels of viral RNA than FR and Delta in the absence of M cells. In the presence of M cells, the levels of BA.1 RNA increased by ∼40-fold after apical infection (Figure 3A). The enhancing effect of M cells was even more striking after infection of the model from the basolateral side. While only background levels of viral RNA were observed in the absence of M cells, the cultures produced about 3-4 orders of magnitude more FR and Delta virus RNA in their presence (Figure 3A). Altogether, BA.1 was generally strongly attenuated and only produced significant albeit low levels of RNA in the presence of M cells.

**Figure 3.**
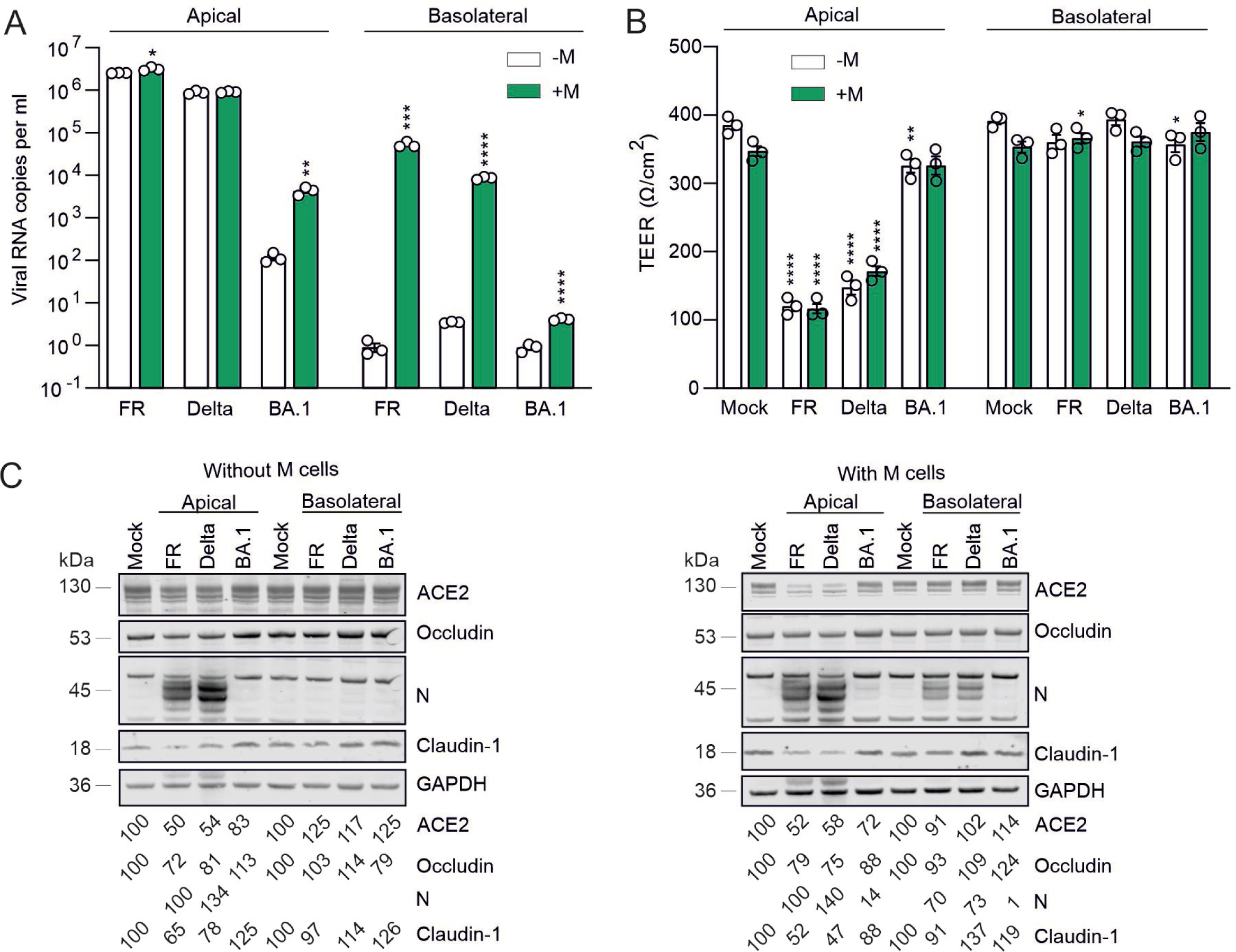
Microfold cells support SARS CoV-2 infection from the basal compartment. (**A**) Analysis of viral RNA copies 48h poi taken from the apical compartments of intestinal epithelial model harboring or lacking M cells, which were infected from the apical or basolateral side. (**B**) TEER and (**C**) WB from A. n=3, the bars represent the mean ± s.e.m.; \**p* = 0.0117, \*\**p* = 0.0016 and \*\*\**p* = 0.0004, **** *p*< 0.0001, unpaired *t*-test.

To assess the effects of the SARS-CoV-2 FR, Delta and BA.1 variants on the integrity of the epithelial monolayer, we determined the TEER values. In agreement with the differences in viral replication, we observed highly significant 2-3-fold increases in permeability after apical inoculation with the FR and Delta strains, while only marginal effects were observed after infection with BA.1 irrespectively of the presence of M cells (Figure 3B, left). In contrast, only modest variations in TEER values were observed after basolateral infection of the intestinal epithelium model despite high replication rates (Figure 3B, right). Western blot analysis confirmed efficient expression of the viral N protein and reduced expression of ACE2, as well as the tight junction proteins claudin-1 and occludin after apical infection with the FR and Delta but not BA.1 SARS-CoV-2 strains (Figures 3C, S2). There was significant FR and Delta N protein expression after basolateral infection in the presence of M cells but no significant changes in the expression levels of tight junction proteins (Figures 3C, S2). This agrees with the results on viral replication and monolayer permeability. Altogether, these results showed that FR and Delta replicate with higher efficiency and damage the gut mucosa more severely than BA.1. M cells promote SARS-CoV-2 replication after basolateral infection but the levels were too low to impair barrier function.

### Remdesivir and Camostat prevent SARS-CoV-2 replication and mucosal damage

To obtain insights into the pathways allowing different SARS-CoV-2 strains to replicate in intestinal epithelial cells, we performed the infection in the presence of the TMPRSS2 inhibitor Camostat and E64d (Hoffmann et al., 2021) inhibiting cathepsins, which may allow processing and cleavage of Spike proteins in endosomes to allow viral entry (Zhao et al., 2021). The fusion inhibitor EK1 (Xia et al., 2022, 2021) and Remdesivir, which inhibits RNA-dependent RNA polymerase (Gordon et al., 2020), served as controls. Remdesivir completely prevented replication of all three SARS-CoV-2 strains (Figure 4A). In comparison, EK1 was highly effective against Delta but displayed little inhibition of the early French (FR) strain, and BA.1 showing an intermediate susceptibility. Camostat inhibited replication of Delta and BA.1 by >95% but only moderately reduced replication (∼30%) of FR even at the highest concentration (Figure 4A). E64d inhibited the Delta strain in a dose dependent manner by up to 70%. However, E64d was inactive against FR and BA.1 and even moderately increased viral RNA production at the lowest (20 µM) concentration. The reason for this is most likely that E64d inhibits autophagy (Yang et al., 2013), which has been reported to restrict SARS-CoV-2 infection (Hayn et al., 2021; Kratzel et al., 2021). In contrast, combinations of Camostat and E64d were generally highly effective (Figure 4A). Altogether, the results indicate that early FR strain uses both TMPRSS2 and Cathepsin dependent entry pathways for efficient replication in this co-culture intestinal model, while Delta and BA.1 are largely dependent on TMPRSS2.

**Figure 4.**
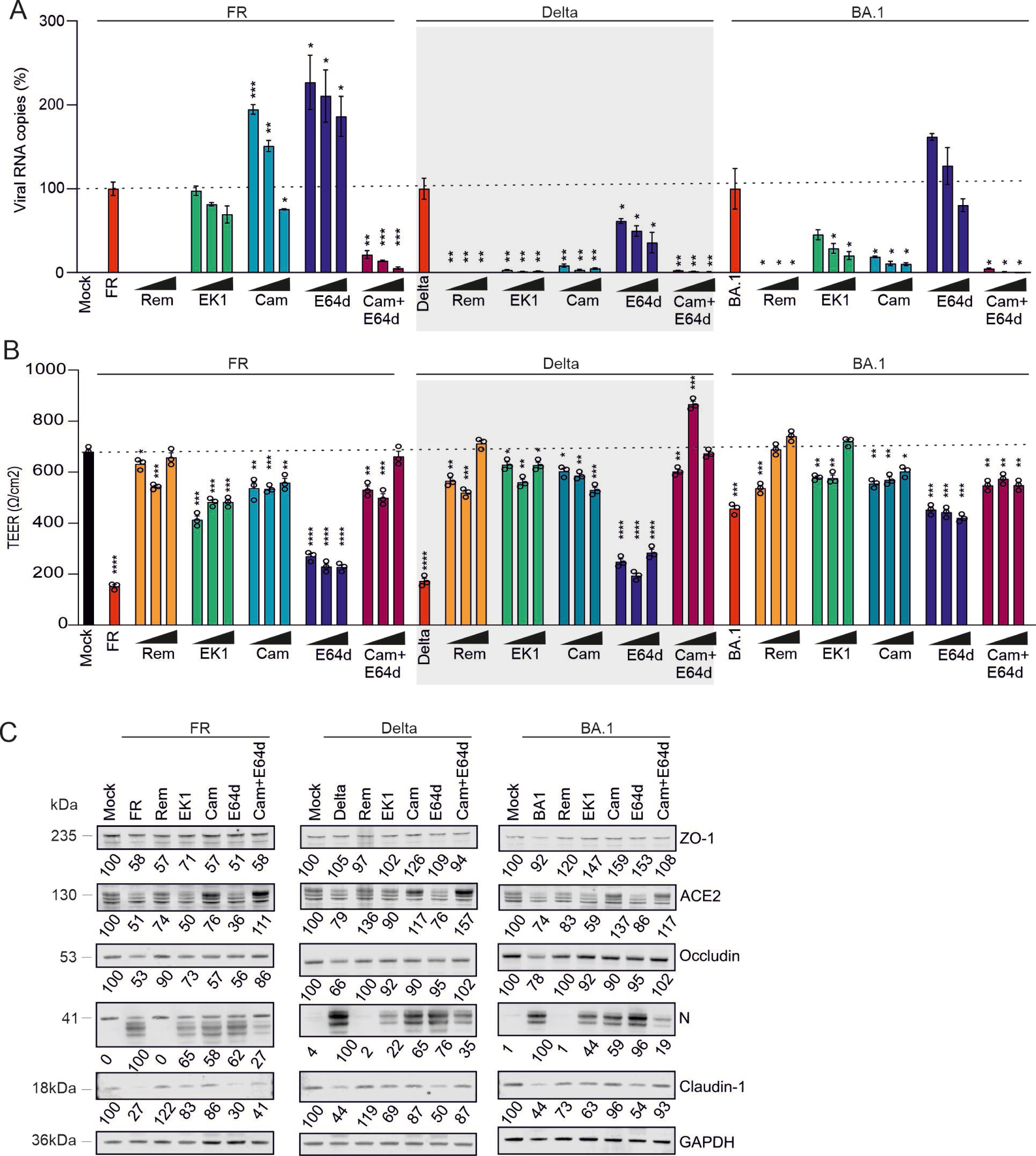
Dependency on different entry and post-entry pathways and tight junction destruction. (**A**) Viral RNA copy numbers, (**B**) TEER values and (**D**) WB of intestinal epithelial model which was before infection with different viral strains of SARS CoV-2 pre-treated with Remdesivir, EK1, Camostat, E64d and combination of E64d and Camostat in increasing concentrations (Remdesivir 5µm, 10 µm and 20 µm; EK1, Camostat, E64d 20 µm, 40 µm and 80 µm). Samples for WB were treated with the highest inhibitors concentration. Inhibitors were freshly added after infection and 24h poi. Samples and measurements were taken 48h poi. n=3, the bars represent the mean ± s.e.m.; \**p* = 0.0117, \*\**p* = 0.0016 and \*\*\**p* = 0.0004, **** *p*< 0.0001, unpaired *t*-test.

In the absence of inhibitors, FR and Delta reduced the TEER values by ∼5-fold, while BA.1 only caused a 1.6-fold reduction (Figure 4B). This decrease in epithelial integrity coincided with decreases of tight junction protein levels. All virus variants downregulated occludin (to 53% FR, 66% Delta and 78% BA.1) and more drastically claudin-1 (to 27% FR, 44% Delta and 44% BA.1; Figure 4C). In contrast, marked decreases of Zonula occludens-1 (ZO-1, also known as Tight junction protein-1) were only observed after infection with the FR strain (Figure 4C). In agreement with their effects on viral replication, Remdesivir, EK1 and the combination of Camostat and E64d prevented disruptive effects on epithelial integrity, while E64d had little protective effect (Figure 4B). All three SARS-CoV-2 strains downregulated ACE2 expression (to 52% FR, 79% Delta and 74% BA.1). Remdesivir, as well as the combination of Camostat and E64d, prevented this reduction. Treatment with Camostat (but not E64d and EK1) alone also prevented loss of ACE2 expression. Altogether, the results indicate that SARS-CoV-2 impairs gut barrier function and the expression of tight junction proteins and further show that Remdesivir, Camostat and (to a lesser extent) EK1 prevent viral replication and associated damaging effects in the co-culture intestinal barrier model.

### Attenuated replication and barrier disruption by SARS-CoV-2 Omicron

Our results showed that BA.1 replicates with lower efficiency and causes less damage to the gut mucosa than the earlier SARS-CoV-2 FR and Delta variants. This agrees with published data reporting that BA.1 is attenuated and less pathogenic compared to early virus strains (Nchioua et al., 2023, 2022; van Doremalen et al., 2022). However, it has been established that subsequent Omicron variants, such as BA.2, the resulting BA.5 VOC, and the wide-spread XBB1.5 variant show increasing replication fitness and possibly also pathogenicity compared to BA.1 (Hoffmann et al., 2023b, 2023a; Kimura et al., 2022; Pastorio et al., 2023; Tamura et al., 2023; Uraki et al., 2022). We found that BA.1 produced moderately lower levels of viral RNA in infected cultures compared to the FR, NL and Delta strains (Figure 5A). The lowest quantities of viral RNA were detected in the supernatant of BA.2-infected model intestinal epithelia and gradual increased over BA.5 to XBB1.5. Altogether, however, the quantities of cell-free viral RNA differed only moderately between the seven different SARS-CoV-2 variants. In contrast, the levels of cell-associated viral RNA of all four Omicron variants were ∼10- to 20-fold lower compared to those detected in model epithelia infected with the FR, NL and Delta strains (Figure 5B). On average, the ratios of cell-free to cell-associated viral RNA were ∼6- to 15-fold higher for the Omicron variants compared to the early FR strain (Figure 5C) indicating efficient virion release. Most notably, all three early SARS-CoV-2 strains significantly increased gut permeability, while the Omicron subvariants had little disruptive effects that only reached significance for XBB1.5 (Figure 5D). Both viral replication as well as inflammatory cytokines may play a role in impaired gut barrier function in COVID-19 patients. However, we detected only modest induction of IFN-stimulated genes (ISGs), such as OAS1 and ISG15, and no significant differences between the seven SARS-CoV-2 variants used (Figure 5E, 5F) suggesting that decreased barrier function was a direct consequence of virus infection.

**Figure 5.**
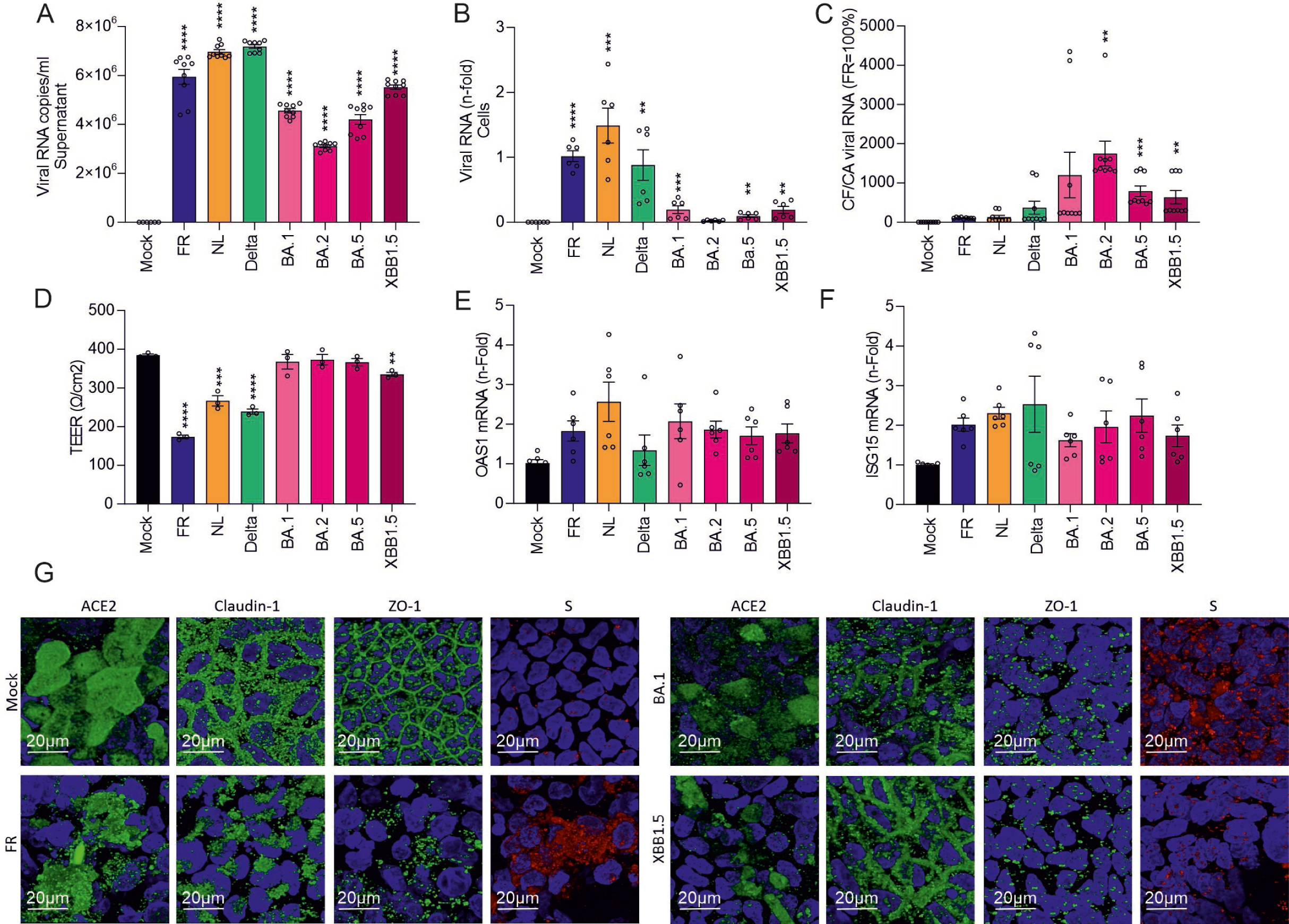
Viral release, epithelial integrity and immune activation of different SARS CoV-2 viral strains. **(A)** Analysis of viral RNA copy numbers in supernatant-cell free (CF) and (**B**) in the cells-cell associated (CA), (**C**) viral release calculated as division of CF with CA viral RNA copies, FR was set to 100%. (**D**) TEER measurements, (**E**) OAS1 and (**F**) ISG15 mRNA levels in intestinal epithelial model infected with different SARS CoV-2 viral strains. n=3-9, the bars represent the mean ± s.e.m.; \**p* = 0.0117, \*\**p* = 0.0016 and \*\*\**p* = 0.0004, **** *p*< 0.0001, unpaired *t*-test. (**G**) Immunofluorescent staining of ACE2, Claudin-1, ZO-1 and S in intestinal epithelial model infected with Mock, FR, BA.1 or XBB1.5 SARS CoV-2 viral strains.

Confocal microscopy confirmed that SARS-CoV-2 FR infection severely affects the expression and localization of claudin-1 and ZO-1 in the co-culture epithelial model (Figure 5G). Both proteins localized mainly in the tight junctions between cells in uninfected epithelial layers. The lattice-like appearance was disturbed and claudin-1 and ZO-1 were only detected in clump or punctate structures in FR infected model epithelia. Infection by BA.1 and XBB1.5 had less drastic effects on the localization of claudin-1. Unexpectedly, however, ZO-1 was also efficiently depleted from tight junctions in epithelial co-cultures infected with the Omicron variants (Figure 5G). It has been reported that ZO-1 is dispensable for barrier function but essential for effective mucosal repair (Kuo et al., 2021). Our finding that Omicron variants deplete ZO-1 from tight junctions but do not increase epithelial permeability agree with this but further suggest that they may predispose the intestinal barrier for damage by preventing efficient repair.

## DISCUSSION

In the present study, we show that co-cultures of epithelial Caco-2 cells with mucus-secreting HT29 goblet cells express high levels of ACE2 and TMPRSS2 and present a useful model to examine interactions between SARS-CoV-2 and the intestinal epithelial barrier. Co-culture with human Raji cells induced differentiation of Caco-2 cells into M-like cells and substantially increased the susceptibility of the model epithelium to basolateral SARS-CoV-2 infection. Early strains of SARS-CoV-2 and the Delta variant exhibited high replication efficiency in this intestinal model, depleted the tight junction proteins claudin-1, occludine as well as ZO-1, and severely impaired mucosal barrier integrity. In comparison, infection by Omicron subvariants had less damaging effects on barrier function but also depleted ZO-1 from tight junctions, although the overall ZO-1 expression levels detected by western blot were hardly affected. Disruption of mucosal integrity was most likely a direct consequence of viral replication since we observed only modest induction of ISGs (OAS1 and ISG15) that did not differ significantly between the different SARS-CoV-2 strains. Altogether, co-cultures of Caco-2 colon epithelial cells, HT29-MTX intestinal mucous-producing goblet cells, and Raji B lymphocytes allow to assess the pathological impact of SARS-CoV-2 on the integrity of the gut epithelial barrier and reveal variant-specific differences that may have implications for disease severity and transmission efficiencies.

Previous studies on SARS-CoV-2 replication and damaging effects in the gut have been performed in gut organoids and small animal models (Frappart et al., 2020; Han et al., 2022; Krüger et al., 2020; Miyakawa et al., 2022; Song et al., 2021). One advantage of the present model is that it includes both human epithelial Caco-2 cells, as well as mucus-secreting HT29 cells, and faithfully replicates the physiological architecture of the human intestinal barrier. The presence of human Raji cells inducing differentiation of Caco-2 cells into M-like cells, which are known to be susceptible to virus infection (Khreefa et al., 2023), further enhances the model’s suitability to study SARS-CoV-2 infection and translocation at the mucosal surface. While organoids showed donor-dependent variations in the SARS-CoV-2 infection (Jang et al., 2022), the present co-culture model was highly susceptible to viral replication and allowed to determine effects on intestinal barrier function, differential effects of SARS-CoV-2 variants, and analysis of antiviral drugs and therapeutic agents. In comparison to small animal models, the triple co-culture model avoids ethical concerns, is more affordable and faster to perform. In addition, it offers the possibility to modulate specific components, such as the ratios of specific cell types, is scalable and well reproducible.

We found that co-culture with Raji cells not only induced differentiation of Caco-2 cells to M-like cells but also substantially increased the susceptibility of the model epithelium to basolateral SARS-CoV-2 infection. M cells are specialized epithelial cells found in the mucosa-associated lymphoid tissue (MALT), that play a role in the mucosal immune response by transporting antigens, including pathogens, from the gut lumen to underlying immune cells (Corr et al., 2008; Mabbott et al., 2013). M-like cells might be particularly susceptible to SARS-CoV-2 (Khreefa et al., 2023) or take up virions and transport them through the barrier (Kimura, 2018). Our results clearly show that M-like cells increase the susceptibility of the intestinal epithelium to productive basolateral SARS-CoV-2 infection suggesting that they help the virus to cross the intestinal epithelial barrier. Further studies are required to clarify the underlying mechanisms and potential implications for viral pathogenesis.

It has been reported that COVID-19 patients suffering from gastrointestinal symptoms show prolonged disease duration and severity of disease (Xu et al., 2023). Thus, drugs suppressing intestinal SARS-CoV-2 replication and damaging effects are of high interest. In agreement to previous results obtained in gut organoids (Krüger et al., 2020), we found that Remdesivir efficiently inhibits SARS-CoV-2 infection and barrier damage in the present epithelial model. The fusion inhibitor EK1 also displayed significant inhibitory and protective effects. EK1 inhibited the Delta variant more efficiently than the early FR SARS-CoV-2 strain. This was unexpected since previous studies have shown that EK1 binds the HR1 domains of numerous highly diverse coronaviruses including SARS-CoV-2 VOCs, MERS-CoV and common cold CoVs (Xia et al., 2022, 2021). Thus, the reasons for the different efficiency of EK1 against FR and Delta need further study. Combined treatment with Camostat and E64d entirely prevented replication and damaging effects of all three SARS-CoV-2 variants investigated, while treatment with E64d alone had little inhibitory effect. Efficient inhibition of Delta and BA.1 by Camostat indicates that these variants are mainly dependent on TMPRSS2 for replication in the intestinal epithelium. In comparison, efficient decrease of FR replication required inhibition of both TMPRSS2 and cathepsins. This came as surprise since previous data suggest that Omicron evolved to become more independent of TMPRSS2 (Meng et al., 2022). Further analyses are required to fully elucidate potential differences in entry pathways. In either case, our results show that treatment with remdesivir might efficiently prevent SARS-CoV-2 replication and gastrointestinal damage and symptoms.

All four Omicron variants investigated showed lower levels of replication compared to earlier SARS-CoV-2 strains and had little if any effects on gut barrier function assessed by TEER values. These results agree with clinical studies showing that gastrointestinal symptoms are relatively rare in Omicron infected individuals (Menni et al., 2022). Direct comparison of the viral RNA levels in the cells and in the supernatants revealed about 5- to 15-fold higher ratios of cell-free to cell-associated vRNA for the four Omicron compared to the early FR and Delta variants. This suggests that Omicron might spread less efficiently in the intestinal epithelium but be released with higher efficiency. This agrees with recent evidence that changes in Omicron Spike increase its ability to counteract tetherin (Shi et al., 2024). These features might contribute to the efficient spread and reduced pathogenicity of Omicron variants.

Reduced replication of Omicron subvariants is consistent with previous results obtained in organoid models (Miyakawa et al., 2022). In agreement with lower infection rates and largely maintained epithelial barrier function, BA.1 and XBB1.5 affected the localization of the tight junction protein claudin-1 less severely than the early FR strain. In striking contrast, ZO-1, a marker of structural and functional integrity was efficiently depleted from tight junctions by all SARS-CoV-2 strains. It has been reported that ZO-1 is dispensable for gut barrier function but critical for effective mucosal repair (Kuo et al., 2021). Thus, impairment of ZO-1 by SARS-CoV-2 infection may prevent restoration of the mucosal barrier and hence contribute to the long-term consequences and complications observed in COVID-19 patients (Xu et al., 2023).

In summary, our study establishes a robust model for SARS-CoV-2 infection in the intestinal epithelium and reveals differential impacts of viral variants on mucosal integrity. Our results add to the evidence that the gut epithelium is highly susceptible to SARS-CoV-2 infection and agrees with clinical findings that COVID-19 is frequently associated with impaired gut barrier function. Positive aspects are that currently dominating Omicron variants seem to be significantly less damaging to the gut mucosa than early SARS-CoV-2 strains. In addition, our results suggest that treatment with Remdesivir and other therapeutic agents will protect the gut mucosa against SARS-CoV-2 infection and associated damage. Since gastrointestinal complications and virus shedding may persist long after the virus is cleared from the respiratory system, further studies on the underlying mechanisms, its role in long-COVID and how to prevent it are warranted.

## MATHERIAL AND METHODS

### Viruses

The SARS-CoV-2 variant B.1.1.529 BA.5 (Omicron BA.5) was kindly provided by Prof. Dr. Florian Schmidt and Dr. Bianca Schulte (University of Bonn). The BetaCoV/ Netherlands/01/NL/2020 (NL-02-2020) lineage, the BetaCoV/ France/IDF0372/2020 (FR, French) and the hCoV-19/Netherlands/NH-EMC-1720/2021 lineage B.1.1.529 (Omicron BA.1) were obtained from the European Virus Archive. The SARS-CoV-2 isolate of lineage B.1.617.2 (Delta) was kindly provided by Prof. Hendrik Streeck, Bonn University Medical Center, Bonn, Germany. The SARS-CoV-2 hCoV-19/USA/CO-CDPHE-2102544747/2021, lineage B.1.1.529, BA.2 (Omicron BA.2) was obtained from the BEI database. The SARS-CoV-2 variant XBB.1.5 (Omicron XBB.1.5) was kindly provided by Viviana Simon (Icahn School of Medicine at Mount Sinai, New York, USA). For propagation of SARS-CoV-2 strains, Vero E6 cells overexpressing ACE2 and TMPRSS2 were seeded to 70-90% confluency in 75 cm² cell culture flasks and inoculated with the SARS-CoV-2 isolates (multiplicity of infection (MOI) of 0.03-0.1) in 3.5 ml serum-free medium. The cells were incubated for 2h at 37°C, before adding 20 ml medium containing 15 mM HEPES. Virus stocks were harvested as soon as strong cytopathic effect (CPE) became apparent. The virus stocks were centrifuged for 5 min at 1,000 g to remove cellular debris, aliquoted, and stored at −80°C until further use. All procedures and assays involving genuine SARS-CoV-2 were performed in the BSL3 facilities at the University of Ulm in accordance with institutional biosafety committee guidelines.

### Cell culture

A human colon adenocarcinoma cell line Caco-2 and HT29-MTX-E12 cells were purchased from ECACC (# 86010202, #12040401) and cultured in DMEM (Gibco) containing 20% fetal bovine serum (Gibco), 1% MEM non-essential amino acid solution (Gibco), 1% L-Glutamine (PAN) and 1% penicillin–streptomycin solution (PAN) under a humidified atmosphere of 5% CO_2_ at 37 °C. Raji cells (ECACC, #85011429) were grown in RPMI medium (Gibco) containing 10% fetal bovine serum (Gibco), 1% L-Glutamine (PAN) and 1% penicillin–streptomycin solution (PAN) under a humidified atmosphere of 5% CO_2_ at 37 °C.

### Intestinal epithelial model

For the development of intestinal epithelial model, 50.000 Caco-2 cells alone or Caco-2 and HT29 cells in ratio 9:1 or 7:3 were seeded on to the TC-Inserts (Sarstedt, #83.3931). Medium was changed every second day. On the day 14, 0.5M Raji cells were added into the lower compartment. Cells were cultivated for another 5 days were Raji density was monitored and keep at 0.5M each day.

### Virus infection

The medium was refreshed and gut models were infected either from above or below the filter with SARS-CoV-2 variants (MOI=0.1). Two hundred microliters of the inoculum were used to infect the cells. Six hours later, input virus was removed and cells were washed with PBS and supplemented with fresh medium. For analysis of infection, supernatants and cells were harvested for qRT-PCR analysis either 24 or 48 hours post-infection. For Western Blot analysis, cells were detached and lysed with 150 microliters of transmembrane lysis buffer. For immunofluorescence, the cells were washed with PBS and fixed in 4% PFA in PBS for 30 minutes.

### TEER measurements

Medium was removed, substituted with the medium kept at room temperature, electrode was inserted and TEER values were measured with the use of World precision instruments, EVOM3.

### Alkaline phosphatase activity

In order to analyze phosphatase activity SIGMAFAST^TM^ (N2770, Sigma) kit was used under the manufacturer instructions.

### Alcian blue staining

Determination of enterocytes/goblet cell ratio and mucin production was performed with Alcian Blue (Sigma Aldrich) staining. Here, cells were placed in 3% acetic acid for 3 min and stained with Alcian Blue solution (1 g of Alcian Blue 8GX in 100 mL of 3% acetic acid) for 30 min. Afterward, cells were rinsed twice with distilled water and visualized by light microscopy.

### Immunostaining

Cells were washed 1x with PBS, fixed with 4% PFA for 20 min, washed 3x with 1x PBS and permeabilized/blocked with 0.5 % Triton-PBS 5% BSA for 2h on RT. Afterwards primary antibodies in 1% BSA were added and incubated on 4°C overnight. Next day cells were washed 3x PBS, secondary antibodies or Phalloidin 647 (Alexa Fluor, Invitrogen; 0.5ul 400x) with DAPI (1/1000) were added for 1h 37°C. Afterwards cells were washed 5x PBS, membranes were cut out of the inserts and imbedded into Vectashield mounting medium onto the microscopy glass slides. Following antibodies were used: ACE2 (ab166755, Abcam), Caludin-1 (ab242370, Abcam), SARS Spike CoV-2 (SARS-CoV / SARS-CoV-2 (COVID-19) spike antibody [1A9], GTX-GTX632604), ZO-1 (ab221547, Abcam), Donkey anti-Rabbit IgG (H+L) Secondary Antibody, Alexa Fluor™ 488 and 647 (A21206, A31573, Invitrogen).

### Scanning electron microscopy

Cells were prepared for SEM by critical point drying as described previously(Schütz et al., 2021). In brief, cells on the transwell membrane/ TC-Inserts were fixed with a fixative containing 2.5% glutaraldehyde, and 1% saccharose in 0.1M phosphate buffer pH 7.3 overnight at 4°C. Post fixation of cells was conducted with 2% osmium tetroxide in PBS for 20 minutes. For dehydration in increasing concentration of isopropanol the filter membrane was cut out of the TC inserts with a scalpel and put into metal containers. In those containers, also critical point drying was conducted. After 2 nm platinum coating by electron beam evaporation, the samples were imaged in a Hitachi S-5200 field emission scanning electron microscope with 10 kV accelerating voltage using the secondary electron signal.

### Western blotting and antibodies

To determine the expression of cellular and viral proteins, medium was removed, cells were washed with 1xPBS and 150ul of Western blot lysis buffer (150 mM NaCl, 50 mM HEPES, 5 mM EDTA, 0.1% NP40, 500 μM Na 3VO4, 500 μM NaF, pH 7.5) supplemented with protease inhibitor (1:500, Roche) was added onto the cells. Cells were incubated with lysis buffer for 15min when they were re-suspended and transferred into 1.5ml tubes. Samples were centrifuged (4 °C, 20 min, 20,817 × g) to remove cell debris, supernatants were transferred to fresh tubes and protein concentrations were determined with BCA Protein Assay Kit (23227, Thermo). Protein concentrations were normalized, samples were mixed with Protein sample loading buffer (Li-COR) with 10% β-mercaptoethanol, heated at 95 °C for 5 min and 40 μg protein was loaded onto 8–15% SDS–PAGE gels. The electrophoresed protein samples were blotted onto Immobilon-FL PVDF (Merck Millipore) membranes. The following antibodies were used: ACE2 (ab166755, Abcam), Claudin-1 (ab242370, Abcam), GAPDH (607902, BioLegend), Occludin (MAB7074, R&D), SARS-CoV-2 (COVID-19) Nucleocapsid antibody [6H3] (GTX632269, GeneTex), TMPRSS2 (ab109131, Abcam).

### qPCR

SARS CoV-2 nucleoprotein (N) RNA levels were determined in supernatants of cells collected from SARS-CoV-2 infected cells 48h post-infection. Total RNA was isolated using the Viral RNA Mini Kit (Qiagen) according to the manufacturer’s instructions. qRT-PCR was performed according to the manufacturer’s instructions using TaqMan Fast Virus 1-Step Master Mix (Thermo Fisher) and an OneStepPlus Real-Time PCR System (96-well format, fast mode). Primers were purchased from Biomers and dissolved in RNAse free water. Synthetic SARS-CoV-2-RNA (Twist Bioscience) were used as a quantitative standard to obtain viral copy numbers. All reactions were run in triplicates using TaqMan primers/probes.

### Detection of viral particles on SEM images

Viral particles have been manually selected, labelled and counted with the help of the program 3dmod from the imod package (Kremer et al., 1996). Our criteria for viral particles were the size (70 to 100 nm, considering that the particle slightly shrink during standard sample preparation for SEM) and the not perfectly round shape with small spike-like protrusions. Since these kinds of particles can hardly be found on the mock-cells, we conclude that the majority of the selected particles are indeed corona viral particles.

### Statistics and reproducibility

Statistical analyses were performed using Graph-Pad PRISM 8 (GraphPad Software). p Values were determined using unpaired *t*-test. Significant differences are denoted by *p <0.05, **p <0.01, ***p <0.001 and ****p <0.0001. Number of independent replicates (n) is indicated for each dataset.

## ACKNOWLEDGEMENTS

We thank Prof. Konstantin Sparrer (Ulm University) for critical reading of the manuscript and helpful discussions, as well as Prof. Florian Schmidt, Dr. Bianca Schulte and Prof. Hendrik Streeck (all University of Bonn, Germany) and Prof. Viviana Simon (Mount Sinai, New York) for providing SARS-CoV-2 variants. This work was supported by the Deutsche Forschungsgemeinschaft (DFG) under CRC1279.

## DATA AVAILABILITY STATEMENT

The data that support the findings of this study are openly available in Mendeley Data; DOI: 10.17632/n2cyh3pk4h.1.

## EXPANDED VIEW FIGURE LEGENDS

**Figure EV1. SEM images of intestinal model infected with FR and Delta SARS CoV-2 strains.** Images are matching images in Figure 2 but without coloring the viral particles.

**Figure EV2. Evaluation of Occludin and Claudin-1 protein expression.** Intestinal model with or without M cells was infected with FR or Delta SARS CoV-2 strains from the apical or basolateral side. Cells were taken for the WB analysis 48h poi. Graph presents evaluation of protein expression from 3 independent experiments where Mock levels were set to 100%. n=3, the bars represent the mean ± s.e.m.; *p = 0.0117, **p = 0.0016 and **** p< 0.0001, unpaired t-test.

## REFERENCES

Ahn, J.H., Kim, J., Hong, S.P., Choi, S.Y., Yang, M.J., Ju, Y.S., Kim, Y.T., Kim, H.M., Rahman, M.T., Chung, M.K., Hong, S.D., Bae, H., Lee, C.-S., Koh, G.Y., 2021. Nasal ciliated cells are primary targets for SARS-CoV-2 replication in the early stage of COVID-19. J Clin Invest 131, e148517. 10.1172/JCI148517

Al-Momani, H., Aolymat, I., Almasri, M., Mahmoud, S.A., Mashal, S., n.d. Prevalence of gastro-intestinal symptoms among COVID-19 patients and the association with disease clinical outcomes. Future Sci OA 9, FSO858. 10.2144/fsoa-2023-0040

Assimakopoulos, S.F., Eleftheriotis, G., Lagadinou, M., Karamouzos, V., Dousdampanis, P., Siakallis, G., Marangos, M., 2022. SARS CoV-2-Induced Viral Sepsis: The Role of Gut Barrier Dysfunction. Microorganisms 10, 1050. 10.3390/microorganisms10051050

Béduneau, A., Tempesta, C., Fimbel, S., Pellequer, Y., Jannin, V., Demarne, F., Lamprecht, A., 2014. A tunable Caco-2/HT29-MTX co-culture model mimicking variable permeabilities of the human intestine obtained by an original seeding procedure. European Journal of Pharmaceutics and Biopharmaceutics 87, 290–298. 10.1016/j.ejpb.2014.03.017

Cabellos, J., Delpivo, C., Fernández-Rosas, E., Vázquez-Campos, S., Janer, G., 2017. Contribution of M-cells and other experimental variables in the translocation of TiO2 nanoparticles across in vitro intestinal models. NanoImpact 5, 51–60. 10.1016/j.impact.2016.12.005

Cappell, M.S., Friedel, D.M., 2023. Gastrointestinal Bleeding in COVID-19-Infected Patients. Gastroenterol Clin North Am 52, 77–102. 10.1016/j.gtc.2022.10.004

Casteleyn, C., Van den Broeck, W., Gebert, A., Tambuyzer, B.R., Van Cruchten, S., Van Ginneken, C., 2013. M cell specific markers in man and domestic animals: valuable tools in vaccine development. Comp Immunol Microbiol Infect Dis 36, 353–364. 10.1016/j.cimid.2013.03.002

Clerbaux, L.-A., Mayasich, S.A., Muñoz, A., Soares, H., Petrillo, M., Albertini, M.C., Lanthier, N., Grenga, L., Amorim, M.-J., 2022. Gut as an Alternative Entry Route for SARS-CoV-2: Current Evidence and Uncertainties of Productive Enteric Infection in COVID-19. Journal of Clinical Medicine 11, 5691. 10.3390/jcm11195691

Corr, S.C., Gahan, C.C.G.M., Hill, C., 2008. M-cells: origin, morphology and role in mucosal immunity and microbial pathogenesis. FEMS Immunology & Medical Microbiology 52, 2–12. 10.1111/j.1574-695X.2007.00359.x

Eleftheriotis, G., Tsounis, E.P., Aggeletopoulou, I., Dousdampanis, P., Triantos, C., Mouzaki, A., Marangos, M., Assimakopoulos, S.F., 2023. Alterations in gut immunological barrier in SARS-CoV-2 infection and their prognostic potential. Frontiers in Immunology 14.

Farsi, Y., Tahvildari, A., Arbabi, M., Vazife, F., Sechi, L.A., Shahidi Bonjar, A.H., Jamshidi, P., Nasiri, M.J., Mirsaeidi, M., 2022. Diagnostic, Prognostic, and Therapeutic Roles of Gut Microbiota in COVID-19: A Comprehensive Systematic Review. Front Cell Infect Microbiol 12, 804644. 10.3389/fcimb.2022.804644

Frappart, P.-O., Walter, K., Gout, J., Beutel, A.K., Morawe, M., Arnold, F., Breunig, M., Barth, T.F., Marienfeld, R., Schulte, L., Ettrich, T., Hackert, T., Svinarenko, M., Rösler, R., Wiese, S., Wiese, H., Perkhofer, L., Müller, M., Lechel, A., Sainz, B., Hermann, P.C., Seufferlein, T., Kleger, A., 2020. Pancreatic cancer-derived organoids - a disease modeling tool to predict drug response. United European gastroenterology journal 8, 594–606. 10.1177/2050640620905183

Gagnon, M., Zihler Berner, A., Chervet, N., Chassard, C., Lacroix, C., 2013. Comparison of the Caco-2, HT-29 and the mucus-secreting HT29-MTX intestinal cell models to investigate Salmonella adhesion and invasion. J Microbiol Methods 94, 274–279. 10.1016/j.mimet.2013.06.027

Gavriatopoulou, M., Korompoki, E., Fotiou, D., Ntanasis-Stathopoulos, I., Psaltopoulou, T., Kastritis, E., Terpos, E., Dimopoulos, M.A., 2020. Organ-specific manifestations of COVID-19 infection. Clin Exp Med 20, 493–506. 10.1007/s10238-020-00648-x

Gordon, C.J., Tchesnokov, E.P., Woolner, E., Perry, J.K., Feng, J.Y., Porter, D.P., Götte, M., 2020. Remdesivir is a direct-acting antiviral that inhibits RNA-dependent RNA polymerase from severe acute respiratory syndrome coronavirus 2 with high potency. J Biol Chem 295, 6785–6797. 10.1074/jbc.RA120.013679

Guimarães Sousa, S., Kleiton de Sousa, A., Maria Carvalho Pereira, C., Sofia Miranda Loiola Araújo, A., de Aguiar Magalhães, D., Vieira de Brito, T., Barbosa, A.L.D.R., 2022. SARS-CoV-2 infection causes intestinal cell damage: Role of interferon’s imbalance. Cytokine 152, 155826. 10.1016/j.cyto.2022.155826

Guo, M., Tao, W., Flavell, R.A., Zhu, S., 2021. Potential intestinal infection and faecal–oral transmission of SARS-CoV-2. Nat Rev Gastroenterol Hepatol 18, 269–283. 10.1038/s41575-021-00416-6

Han, Y., Yang, L., Lacko, L.A., Chen, S., 2022. Human organoid models to study SARS-CoV-2 infection. Nat Methods 19, 418–428. 10.1038/s41592-022-01453-y

Hayashi, Y., Wagatsuma, K., Nojima, M., Yamakawa, T., Ichimiya, T., Yokoyama, Y., Kazama, T., Hirayama, D., Nakase, H., 2021. The characteristics of gastrointestinal symptoms in patients with severe COVID-19: a systematic review and meta-analysis. J Gastroenterol 56, 409–420. 10.1007/s00535-021-01778-z

Hayn, M., Hirschenberger, M., Koepke, L., Nchioua, R., Straub, J.H., Klute, S., Hunszinger, V., Zech, F., Prelli Bozzo, C., Aftab, W., Christensen, M.H., Conzelmann, C., Müller, J.A., Srinivasachar Badarinarayan, S., Stürzel, C.M., Forne, I., Stenger, S., Conzelmann, K.-K., Münch, J., Schmidt, F.I., Sauter, D., Imhof, A., Kirchhoff, F., Sparrer, K.M.J., 2021. Systematic functional analysis of SARS-CoV-2 proteins uncovers viral innate immune antagonists and remaining vulnerabilities. Cell Rep 35, 109126. 10.1016/j.celrep.2021.109126

Hoffmann, M., Arora, P., Nehlmeier, I., Kempf, A., Cossmann, A., Schulz, S.R., Morillas Ramos, G., Manthey, L.A., Jäck, H.-M., Behrens, G.M.N., Pöhlmann, S., 2023a. Profound neutralization evasion and augmented host cell entry are hallmarks of the fast-spreading SARS-CoV-2 lineage XBB.1.5. Cell Mol Immunol 20, 419–422. 10.1038/s41423-023-00988-0

Hoffmann, M., Hofmann-Winkler, H., Smith, J.C., Krüger, N., Arora, P., Sørensen, L.K., Søgaard, O.S., Hasselstrøm, J.B., Winkler, M., Hempel, T., Raich, L., Olsson, S., Danov, O., Jonigk, D., Yamazoe, T., Yamatsuta, K., Mizuno, H., Ludwig, S., Noé, F., Kjolby, M., Braun, A., Sheltzer, J.M., Pöhlmann, S., 2021. Camostat mesylate inhibits SARS-CoV-2 activation by TMPRSS2-related proteases and its metabolite GBPA exerts antiviral activity. EBioMedicine 65, 103255. 10.1016/j.ebiom.2021.103255

Hoffmann, M., Kleine-Weber, H., Schroeder, S., Krüger, N., Herrler, T., Erichsen, S., Schiergens, T.S., Herrler, G., Wu, N.-H., Nitsche, A., Müller, M.A., Drosten, C., Pöhlmann, S., 2020. SARS-CoV-2 Cell Entry Depends on ACE2 and TMPRSS2 and Is Blocked by a Clinically Proven Protease Inhibitor. Cell. 10.1016/j.cell.2020.02.052

Hoffmann, M., Wong, L.-Y.R., Arora, P., Zhang, L., Rocha, C., Odle, A., Nehlmeier, I., Kempf, A., Richter, A., Halwe, N.J., Schön, J., Ulrich, L., Hoffmann, D., Beer, M., Drosten, C., Perlman, S., Pöhlmann, S., 2023b. Omicron subvariant BA.5 efficiently infects lung cells. Nat Commun 14, 3500. 10.1038/s41467-023-39147-4

Jang, K.K., Kaczmarek, M.E., Dallari, S., Chen, Y.-H., Tada, T., Axelrad, J., Landau, N.R., Stapleford, K.A., Cadwell, K., 2022. Variable susceptibility of intestinal organoid– derived monolayers to SARS-CoV-2 infection. PLOS Biology 20, e3001592. 10.1371/journal.pbio.3001592

Jiao, L., Li, H., Xu, J., Yang, M., Ma, C., Li, J., Zhao, S., Wang, H., Yang, Y., Yu, W., Wang, J., Yang, J., Long, H., Gao, J., Ding, K., Wu, D., Kuang, D., Zhao, Y., Liu, J., Lu, S., Liu, H., Peng, X., 2021. The Gastrointestinal Tract Is an Alternative Route for SARS-CoV-2 Infection in a Nonhuman Primate Model. Gastroenterology 160, 1647– 1661. 10.1053/j.gastro.2020.12.001

Jin, B., Singh, R., Ha, S.E., Zogg, H., Park, P.J., Ro, S., 2021. Pathophysiological mechanisms underlying gastrointestinal symptoms in patients with COVID-19. World J Gastroenterol 27, 2341–2352. 10.3748/wjg.v27.i19.2341

Kariyawasam, J.C., Jayarajah, U., Riza, R., Abeysuriya, V., Seneviratne, S.L., 2021. Gastrointestinal manifestations in COVID-19. Trans R Soc Trop Med Hyg 115, 1362–1388. 10.1093/trstmh/trab042

Khreefa, Z., Barbier, M.T., Koksal, A.R., Love, G., Del Valle, L., 2023. Pathogenesis and Mechanisms of SARS-CoV-2 Infection in the Intestine, Liver, and Pancreas. Cells 12, 262. 10.3390/cells12020262

Kim, Y.S., Ho, S.B., 2010. Intestinal Goblet Cells and Mucins in Health and Disease: Recent Insights and Progress. Curr Gastroenterol Rep 12, 319–330. 10.1007/s11894-010-0131-2

Kimura, I., Yamasoba, D., Tamura, T., Nao, N., Oda, Y., Mitoma, S., Ito, J., Nasser, H., Zahradnik, J., Uriu, K., Fujita, S., Kosugi, Y., Wang, L., Tsuda, M., Kishimoto, M., Ito, H., Suzuki, R., Shimizu, R., Begum, M.M., Yoshimatsu, K., Sasaki, J., Sasaki-Tabata, K., Yamamoto, Y., Nagamoto, T., Kanamune, J., Kobiyama, K., Asakura, H., Nagashima, M., Sadamasu, K., Yoshimura, K., Kuramochi, J., Schreiber, G., Ishii, K.J., Hashiguchi, T., Consortium, T.G. to P.J. (G2P-J., Ikeda, T., Saito, A., Fukuhara, T., Tanaka, S., Matsuno, K., Sato, K., 2022. Virological characteristics of the novel SARS-CoV-2 Omicron variants including BA.2.12.1, BA.4 and BA.5. 10.1101/2022.05.26.493539

Kimura, S., 2018. Molecular insights into the mechanisms of M-cell differentiation and transcytosis in the mucosa-associated lymphoid tissues. Anat Sci Int 93, 23–34. 10.1007/s12565-017-0418-6

Kratzel, A., Kelly, J.N., V’kovski, P., Portmann, J., Brüggemann, Y., Todt, D., Ebert, N., Shrestha, N., Plattet, P., Staab-Weijnitz, C.A., von Brunn, A., Steinmann, E., Dijkman, R., Zimmer, G., Pfaender, S., Thiel, V., 2021. A genome-wide CRISPR screen identifies interactors of the autophagy pathway as conserved coronavirus targets. PLoS Biol 19, e3001490. 10.1371/journal.pbio.3001490

Kremer, J.R., Mastronarde, D.N., McIntosh, J.R., 1996. Computer visualization of three-dimensional image data using IMOD. J Struct Biol 116, 71–76. 10.1006/jsbi.1996.0013

Krüger, J., Groß, R., Conzelmann, C., Müller, J.A., Koepke, L., Sparrer, K.M.J., Weil, T., Schütz, D., Seufferlein, T., Barth, T.F.E., Stenger, S., Heller, S., Münch, J., Kleger, A., 2020. Drug Inhibition of SARS-CoV-2 Replication in Human Pluripotent Stem Cell-Derived Intestinal Organoids. Cell Mol Gastroenterol Hepatol. 10.1016/j.jcmgh.2020.11.003

Kuo, W.-T., Zuo, L., Odenwald, M.A., Madha, S., Singh, G., Gurniak, C.B., Abraham, C., Turner, J.R., 2021. The Tight Junction Protein ZO-1 Is Dispensable for Barrier Function but Critical for Effective Mucosal Repair. Gastroenterology 161, 1924– 1939. 10.1053/j.gastro.2021.08.047

Lamers, M.M., Haagmans, B.L., 2022. SARS-CoV-2 pathogenesis. Nat Rev Microbiol 20, 270–284. 10.1038/s41579-022-00713-0

Lee, I.T., Nakayama, T., Wu, C.-T., Goltsev, Y., Jiang, S., Gall, P.A., Liao, C.-K., Shih, L.-C., Schürch, C.M., McIlwain, D.R., Chu, P., Borchard, N.A., Zarabanda, D., Dholakia, S.S., Yang, A., Kim, D., Chen, H., Kanie, T., Lin, C.-D., Tsai, M.-H., Phillips, K.M., Kim, R., Overdevest, J.B., Tyler, M.A., Yan, C.H., Lin, C.-F., Lin, Y.-T., Bau, D.-T., Tsay, G.J., Patel, Z.M., Tsou, Y.-A., Tzankov, A., Matter, M.S., Tai, C.-J., Yeh, T.-H., Hwang, P.H., Nolan, G.P., Nayak, J.V., Jackson, P.K., 2020. ACE2 localizes to the respiratory cilia and is not increased by ACE inhibitors or ARBs. Nat Commun 11, 5453. 10.1038/s41467-020-19145-6

Mabbott, N.A., Donaldson, D.S., Ohno, H., Williams, I.R., Mahajan, A., 2013. Microfold (M) cells: important immunosurveillance posts in the intestinal epithelium. Mucosal Immunol 6, 666–677. 10.1038/mi.2013.30

Mahler, G.J., Shuler, M.L., Glahn, R.P., 2009. Characterization of Caco-2 and HT29-MTX cocultures in an in vitro digestion/cell culture model used to predict iron bioavailability. The Journal of Nutritional Biochemistry 20, 494–502. 10.1016/j.jnutbio.2008.05.006

Masuda, K., Kajikawa, A., Igimi, S., 2011. Establishment and Evaluation of an in vitro M Cell Model using C2BBe1 Cells and Raji Cells. Biosci Microflora 30, 37–44. 10.12938/bifidus.30.37

Meng, B., Abdullahi, A., Ferreira, I.A.T.M., Goonawardane, N., Saito, A., Kimura, I., Yamasoba, D., Gerber, P.P., Fatihi, S., Rathore, S., Zepeda, S.K., Papa, G., Kemp, S.A., Ikeda, T., Toyoda, M., Tan, T.S., Kuramochi, J., Mitsunaga, S., Ueno, T., Shirakawa, K., Takaori-Kondo, A., Brevini, T., Mallery, D.L., Charles, O.J., Bowen, J.E., Joshi, A., Walls, A.C., Jackson, L., Martin, D., Smith, K.G.C., Bradley, J., Briggs, J.A.G., Choi, J., Madissoon, E., Meyer, K.B., Mlcochova, P., Ceron-Gutierrez, L., Doffinger, R., Teichmann, S.A., Fisher, A.J., Pizzuto, M.S., de Marco, A., Corti, D., Hosmillo, M., Lee, J.H., James, L.C., Thukral, L., Veesler, D., Sigal, A., Sampaziotis, F., Goodfellow, I.G., Matheson, N.J., Sato, K., Gupta, R.K., 2022. Altered TMPRSS2 usage by SARS-CoV-2 Omicron impacts infectivity and fusogenicity. Nature 603, 706–714. 10.1038/s41586-022-04474-x

Menni, C., Valdes, A.M., Polidori, L., Antonelli, M., Penamakuri, S., Nogal, A., Louca, P., May, A., Figueiredo, J.C., Hu, C., Molteni, E., Canas, L., Österdahl, M.F., Modat, M., Sudre, C.H., Fox, B., Hammers, A., Wolf, J., Capdevila, J., Chan, A.T., David, S.P., Steves, C.J., Ourselin, S., Spector, T.D., 2022. Symptom prevalence, duration, and risk of hospital admission in individuals infected with SARS-CoV-2 during periods of omicron and delta variant dominance: a prospective observational study from the ZOE COVID Study. The Lancet 399, 1618–1624. 10.1016/S0140-6736(22)00327-0

Miyakawa, K., Machida, M., Kawasaki, T., Nishi, M., Akutsu, H., Ryo, A., 2022. Reduced Replication Efficacy of Severe Acute Respiratory Syndrome Coronavirus 2 Omicron Variant in “Mini-gut” Organoids. Gastroenterology 163, 514–516. 10.1053/j.gastro.2022.04.043

Nchioua, R., Diofano, F., Noettger, S., von Maltitz, P., Stenger, S., Zech, F., Münch, J., Sparrer, K.M.J., Just, S., Kirchhoff, F., 2022. Strong attenuation of SARS-CoV-2 Omicron BA.1 and increased replication of the BA.5 subvariant in human cardiomyocytes. Sig Transduct Target Ther 7, 1–3. 10.1038/s41392-022-01256-9

Nchioua, R., Schundner, A., Klute, S., Koepke, L., Hirschenberger, M., Noettger, S., Fois, G., Zech, F., Graf, A., Krebs, S., Braubach, P., Blum, H., Stenger, S., Kmiec, D., Frick, M., Kirchhoff, F., Sparrer, K.M., 2023. Reduced replication but increased interferon resistance of SARS-CoV-2 Omicron BA.1. Life Sci Alliance 6, e202201745. 10.26508/lsa.202201745

Neuberger, M., Jungbluth, A., Irlbeck, M., Streitparth, F., Burian, M., Kirchner, T., Werner, J., Rudelius, M., Knösel, T., 2022. Duodenal tropism of SARS-CoV-2 and clinical findings in critically ill COVID-19 patients. Infection 50, 1111–1120. 10.1007/s15010-022-01769-z

Pan, F., Han, L., Zhang, Y., Yu, Y., Liu, J., 2015. Optimization of Caco-2 and HT29 co-culture in vitro cell models for permeability studies. International Journal of Food Sciences and Nutrition 66, 680–685. 10.3109/09637486.2015.1077792

Pastorio, C., Noettger, S., Nchioua, R., Zech, F., Sparrer, K.M.J., Kirchhoff, F., 2023. Impact of mutations defining SARS-CoV-2 Omicron subvariants BA.2.12.1 and BA.4/5 on Spike function and neutralization. iScience 26. 10.1016/j.isci.2023.108299

Perrotta, F., Matera, M.G., Cazzola, M., Bianco, A., 2020. Severe respiratory SARS-CoV2 infection: Does ACE2 receptor matter? Respir Med 168, 105996. 10.1016/j.rmed.2020.105996

Rahban, M., Stanek, A., Hooshmand, A., Khamineh, Y., Ahi, S., Kazim, S.N., Ahmad, F., Muronetz, V., Samy Abousenna, M., Zolghadri, S., Saboury, A.A., 2021. Infection of Human Cells by SARS-CoV-2 and Molecular Overview of Gastrointestinal, Neurological, and Hepatic Problems in COVID-19 Patients. Journal of Clinical Medicine 10, 4802. 10.3390/jcm10214802

Scaldaferri, F., Ianiro, G., Privitera, G., Lopetuso, L.R., Vetrone, L.M., Petito, V., Pugliese, D., Neri, M., Cammarota, G., Ringel, Y., Costamagna, G., Gasbarrini, A., Boskoski, I., Armuzzi, A., 2020. The Thrilling Journey of SARS-CoV-2 into the Intestine: From Pathogenesis to Future Clinical Implications. Inflamm Bowel Dis 26, 1306–1314. 10.1093/ibd/izaa181

Schütz, D., Rode, S., Read, C., Müller, J.A., Glocker, B., Sparrer, K.M.J., Fackler, O.T., Walther, P., Münch, J., 2021. Viral Transduction Enhancing Effect of EF-C Peptide Nanofibrils Is Mediated by Cellular Protrusions. Advanced Functional Materials 31, 2104814. 10.1002/adfm.202104814

Shi, Y., Simpson, S., Chen, Y., Aull, H., Benjamin, J., Serra-Moreno, R., 2024. Mutations accumulated in the Spike of SARS-CoV-2 Omicron allow for more efficient counteraction of the restriction factor BST2/Tetherin. PLOS Pathogens 20, e1011912. 10.1371/journal.ppat.1011912

Song, Z., Bao, L., Yu, P., Qi, F., Gong, S., Wang, J., Zhao, B., Liu, M., Han, Y., Deng, W., Liu, J., Wei, Q., Xue, J., Zhao, W., Qin, C., 2021. SARS-CoV-2 Causes a Systemically Multiple Organs Damages and Dissemination in Hamsters. Frontiers in Microbiology 11.

Tamura, T., Ito, J., Uriu, K., Zahradnik, J., Kida, I., Anraku, Y., Nasser, H., Shofa, M., Oda, Y., Lytras, S., Nao, N., Itakura, Y., Deguchi, S., Suzuki, R., Wang, L., Begum, M.M., Kita, S., Yajima, H., Sasaki, J., Sasaki-Tabata, K., Shimizu, R., Tsuda, M., Kosugi, Y., Fujita, S., Pan, L., Sauter, D., Yoshimatsu, K., Suzuki, S., Asakura, H., Nagashima, M., Sadamasu, K., Yoshimura, K., Yamamoto, Y., Nagamoto, T., Schreiber, G., Maenaka, K., Hashiguchi, T., Ikeda, T., Fukuhara, T., Saito, A., Tanaka, S., Matsuno, K., Takayama, K., Sato, K., 2023. Virological characteristics of the SARS-CoV-2 XBB variant derived from recombination of two Omicron subvariants. Nat Commun 14, 2800. 10.1038/s41467-023-38435-3

Tsounis, E.P., Triantos, C., Konstantakis, C., Marangos, M., Assimakopoulos, S.F., 2023. Intestinal barrier dysfunction as a key driver of severe COVID-19. World J Virol 12, 68–90. 10.5501/wjv.v12.i2.68

Uraki, R., Halfmann, P.J., Iida, S., Yamayoshi, S., Furusawa, Y., Kiso, M., Ito, M., Iwatsuki-Horimoto, K., Mine, S., Kuroda, M., Maemura, T., Sakai-Tagawa, Y., Ueki, H., Li, R., Liu, Y., Larson, D., Fukushi, S., Watanabe, S., Maeda, K., Pekosz, A., Kandeil, A., Webby, R.J., Wang, Z., Imai, M., Suzuki, T., Kawaoka, Y., 2022. Characterization of SARS-CoV-2 Omicron BA.4 and BA.5 isolates in rodents. Nature 612, 540–545. 10.1038/s41586-022-05482-7

van Doremalen, N., Singh, M., Saturday, T.A., Yinda, C.K., Perez-Perez, L., Bohler, W.F., Weishampel, Z.A., Lewis, M., Schulz, J.E., Williamson, B.N., Meade-White, K., Gallogly, S., Okumura, A., Feldmann, F., Lovaglio, J., Hanley, P.W., Shaia, C., Feldmann, H., de Wit, E., Munster, V.J., Rosenke, K., 2022. SARS-CoV-2 Omicron BA.1 and BA.2 are attenuated in rhesus macaques as compared to Delta. bioRxiv 2022.08.01.502390. 10.1101/2022.08.01.502390

V’kovski, P., Kratzel, A., Steiner, S., Stalder, H., Thiel, V., 2021. Coronavirus biology and replication: implications for SARS-CoV-2. Nat Rev Microbiol 19, 155–170. 10.1038/s41579-020-00468-6

Xia, S., Chan, J.F.-W., Wang, L., Jiao, F., Chik, K.K.-H., Chu, H., Lan, Q., Xu, W., Wang, Q., Wang, C., Yuen, K.-Y., Lu, L., Jiang, S., 2022. Peptide-based pan-CoV fusion inhibitors maintain high potency against SARS-CoV-2 Omicron variant. Cell Res 32, 404–406. 10.1038/s41422-022-00617-x

Xia, S., Lan, Q., Zhu, Y., Wang, C., Xu, W., Li, Y., Wang, L., Jiao, F., Zhou, J., Hua, C., Wang, Q., Cai, X., Wu, Y., Gao, J., Liu, H., Sun, G., Münch, J., Kirchhoff, F., Yuan, Z., Xie, Y., Sun, F., Jiang, S., Lu, L., 2021. Structural and functional basis for pan-CoV fusion inhibitors against SARS-CoV-2 and its variants with preclinical evaluation. Sig Transduct Target Ther 6, 1–10. 10.1038/s41392-021-00712-2

Xu, E., Xie, Y., Al-Aly, Z., 2023. Long-term gastrointestinal outcomes of COVID-19. Nat Commun 14, 983. 10.1038/s41467-023-36223-7

Yamada, S., Noda, T., Okabe, K., Yanagida, S., Nishida, M., Kanda, Y., 2022. SARS-CoV-2 induces barrier damage and inflammatory responses in the human iPSC-derived intestinal epithelium. Journal of Pharmacological Sciences 149, 139–146. 10.1016/j.jphs.2022.04.010

Yang, Y., Hu, L., Zheng, H., Mao, C., Hu, W., Xiong, K., Wang, F., Liu, C., 2013. Application and interpretation of current autophagy inhibitors and activators. Acta Pharmacol Sin 34, 625–635. 10.1038/aps.2013.5

Zhang, J., Garrett, S., Sun, J., 2021. Gastrointestinal symptoms, pathophysiology, and treatment in COVID-19. Genes Dis 8, 385–400. 10.1016/j.gendis.2020.08.013

Zhao, M.-M., Yang, W.-L., Yang, F.-Y., Zhang, L., Huang, W.-J., Hou, W., Fan, C.-F., Jin, R.-H., Feng, Y.-M., Wang, Y.-C., Yang, J.-K., 2021. Cathepsin L plays a key role in SARS-CoV-2 infection in humans and humanized mice and is a promising target for new drug development. Sig Transduct Target Ther 6, 1–12. 10.1038/s41392-021-00558-8

Zhong, P., Xu, J., Yang, D., Shen, Y., Wang, L., Feng, Y., Du, C., Song, Y., Wu, C., Hu, X., Sun, Y., 2020. COVID-19-associated gastrointestinal and liver injury: clinical features and potential mechanisms. Sig Transduct Target Ther 5, 1–8. 10.1038/s41392-020-00373-7

Zuo, T., Zhang, F., Lui, G.C.Y., Yeoh, Y.K., Li, A.Y.L., Zhan, H., Wan, Y., Chung, A.C.K., Cheung, C.P., Chen, N., Lai, C.K.C., Chen, Z., Tso, E.Y.K., Fung, K.S.C., Chan, V., Ling, L., Joynt, G., Hui, D.S.C., Chan, F.K.L., Chan, P.K.S., Ng, S.C., 2020. Alterations in Gut Microbiota of Patients With COVID-19 During Time of Hospitalization. Gastroenterology 159, 944–955.e8. 10.1053/j.gastro.2020.05.048

